# Induction of X-chromosome Inactivation by the Histone Demethylase SMCX/KDM5C

**DOI:** 10.1101/175174

**Authors:** Srimonta Gayen, Emily Maclary, Yumie Murata-Nakamura, Christina N. Vallianatos, Robert S. Porter, Patricia M. Garay, Shigeki Iwase, Sundeep Kalantry

## Abstract

*XY* male and *XX* female mammals equalize X-linked gene expression through the mitotically-stable transcriptional inactivation of an X-chromosome in females. Although most genes are silent on the inactive-X, some escape silencing and are expressed at higher levels in females vs. males. Here, we show that the escapee *Smcx*/*Kdm5c*, encoding a histone H3K4me2/3 demethylase, underlies the female-specific induction of X-inactivation. Mouse embryonic epiblast cells and differentiating embryonic stem cells (ESCs) lacking SMCX show reduced expression of Xist RNA, which is required for X-inactivation. *Smcx*-heterozygous epiblast cells do not silence X-linked genes efficiently, despite robust *Xist* expression. Overexpression of mouse or human SMCX, but not a catalytically-inactive SMCX or the Y-chromosome homolog SMCY, is sufficient to induce *Xist* and, separately, to silence X-linked genes in male ESCs. Finally, SMCX dose is inversely correlated with H3K4me2 at X-linked loci. Thus, X-inactivation initiates through the evolutionarily conserved, dose-dependent function of the histone demethylase SMCX.

## Main Text

X-inactivation is an evolutionarily conserved process that equalizes X-linked gene expression between male and female mammals by silencing genes on one of the two X-chromosomes in females^1-3^. Once inactivated, replicated copies of the X-chromosome are maintained as inactive throughout future rounds of cell division^3^, thus making X-inactivation a paradigm of epigenetic inheritance. Stable X-inactivation requires the Xist long noncoding RNA^4,5^, which is upregulated from and physically coats the future inactive X-chromosome^6-10^. Xist RNA accumulation recruits proteins that in turn are believed to silence X-linked genes^11-13^. However, the precise order of molecular mechanisms by which female cells upregulate Xist RNA and undergo X-inactivation remain obscure.

We hypothesized that the increased dose in *XX* females compared to *XY* males of one or more X-linked genes that escape X-inactivation induces *Xist* and, separately, silences X-linked genes selectively in females^14^. We thus nominated the X-inactivation escapee *Smcx*/*Kdm5c* as a candidate inducer of *Xist* and X-linked gene silencing, for the following reasons. First, SMCX demethylates histone H3 di-and tri-methylated at lysine 4 (H3K4me2 and H3K4me3)^15,16^, which are chromatin marks associated with active gene expression^17,18^. Second, hypomethylation of H3K4 is associated with *Xist* induction and silencing of X-linked genes on the inactive-X^19,20^. Finally, unlike many other escapees, the *Smcx* gene escapes X-inactivation in both mouse and human^21-24^, suggesting an evolutionarily conserved dose-dependent function.

### ***Smcx* is required to induce Xist RNA**

To test for a role of *Smcx* in X-inactivation, we took advantage of a conditional *Smcx*^fl^ allele to generate homozygous *Smcx*^Δ/Δ^ and hemizygous *Smcx*^Δ/Y^ mice^25^. Strikingly, although *Smcx*^Δ/Y^ males were viable, *Smcx*^Δ/Δ^ females were not (Fig. 1a). We also crossed *Smcx*^fl/Δ^ females harboring a *Zp3-Cre* transgene with *Smcx*^Δ/Y^ males. The expression of the *Zp3-Cre* transgene deletes floxed alleles in growing oocytes and thus ensures that the resulting embryos are depleted of any oocyte-derived maternal SMCX protein^26^, which may influence X-inactivation patterns in the early embryo. This cross also produced *Smcx*^Δ/Y^ males but no *Smcx*^Δ/Δ^ females (Fig. 1a).

To determine if females were perishing due to defects in X-inactivation, we first examined preimplantation mouse embryos. All cells of the female preimplantation mouse embryo undergo imprinted X-inactivation of the paternal X-chromosome^27-29^. We generated *Smcx*^Δ/Δ^ embryonic day (E) 3.5 blastocyst-stage embryos from a cross of *Zp3-Cre*;*Smcx*^fl/Δ^ females with *Smcx*^Δ/Y^ males harboring an X-linked *Gfp* transgene (*X-Gfp*;*Smcx*^Δ/Y^). The paternally-transmitted *X-Gfp* transgene permits visual sexing of the embryos since it’s only transmitted to female progeny^30,31^. We found that *Smcx*^Δ/Δ^ and control *Smcx*^fl/fl^ female E3.5 embryos displayed similar patterns of imprinted Xist RNA induction and X-linked gene silencing by RNA fluorescent *in situ* hybridization (FISH) (Extended Data Fig. 1a).

To delineate when *Smcx*^Δ/Δ^ females were being lost, we next examined E5.5 and E6.5 post-implantation stage embryos. Whereas the control crosses generated the expected distribution of female and male embryos, the test cross yielded significantly fewer E5.5 and E6.5 *Smcx*^Δ/Δ^ female embryos compared to *Smcx*^Δ/Y^ male embryos (Fig. 1b). Consistent with the death of the *Smcx*^Δ/Δ^ female embryos, significantly more embryos were resorbed in the test cross compared to the control crosses. Moreover, the surviving *Smcx*^Δ/Δ^ E5.5 embryos were smaller relative to their heterozygous *Smcx*^fl/Δ^ female and hemizygous *Smcx*^Δ/Y^ male littermates (Extended Data Fig. 1b).

We hypothesized that the under-representation of E5.5 *Smcx*^Δ/Δ^ embryos is due to defective random X-inactivation, which begins between E5.0-E5.25 stage of embryogenesis^32,33^. We therefore examined Xist RNA expression in epiblasts of surviving E5.5 *Smcx*^Δ/Δ^ female embryos. Compared to *Smcx*^fl/fl^ and *Smcx*^fl/Δ^ genotypes, many nuclei in E5.5 *Smcx*^Δ/Δ^ embryonic epiblasts lacked Xist RNA coating by fluorescence *in situ* hybridization (FISH) (Fig. 1c-d). Moreover, many *Smcx*^Δ/Δ^ nuclei with Xist RNA coating exhibited qualitatively weaker Xist RNA foci compared to the control *Smcx*^fl/fl^ and *Smcx*^fl/Δ^ nuclei. To test if the deficiency in Xist RNA coating is due to a failure of differentiation of the pluripotent epiblast progenitor cells, we concurrently assayed expression of the REX1 protein by immunofluorescence (IF). REX1 marks pluripotent epiblast progenitor cells in E3.5 embryos and is rapidly downregulated when these cells differentiate^33^, which is accompanied by the onset of random X-inactivation^33,34^. Unlike E3.5 embryos, the E5.5 control and *Smcx*^Δ/Δ^ epiblasts were devoid of REX1 expression, indicating that they had undergone differentiation (Fig. 1c). In agreement with the RNA FISH data, RT-qPCR also revealed significantly reduced Xist RNA expression but similar *Rex1* RNA levels in E5.5 *Smcx*^Δ/Δ^ compared to *Smcx*^fl/fl^ and *Smcx*^fl/Δ^ epiblasts (Fig. 1e).

To evaluate X-inactivation *in vitro*, we next derived *Smcx*^fl/fl^, *Smcx*^fl/Δ^, and *Smcx*^Δ/Δ^ embryonic stem cells (ESCs). ESCs can be differentiated into epiblast-like cells (EpiLCs)^35^, which mimic the early mouse epiblast just after X-inactivation has initiated^33^. Whereas *Smcx*^fl/fl^ and *Smcx*^fl/Δ^ EpiLCs displayed Xist RNA expression and coating, *Smcx*^Δ/Δ^ EpiLCs were deficient in both (Fig. 1f-h; Extended Data Fig. 1c). Thus, SMCX is required for Xist RNA induction *in vivo* and *in vitro*.

### **Dose-dependent requirement of *Smcx* in X-linked gene silencing**

We next asked if SMCX is required for X-linked gene silencing, by examining heterozygous *Smcx*^fl/Δ^ epiblasts and EpiLCs, which display robust Xist RNA expression and coating (Fig. 1c-h). Due to potential cell-to-cell and gene-to-gene variability in silencing^14^, we employed RNA FISH to compare the expression of eight different genes distributed across the X-chromosome in *Smcx*^fl/fl^ and *Smcx*^fl/Δ^ E5.5 embryonic epiblasts and EpiLCs (Fig. 2a; Extended Data Fig. 2a). A subset of the tested genes, *Lamp2*, *Mecp2*, and *Atrx*, were expressed from the Xist RNA-coated X-chromosome in significantly more *Smcx*^fl/Δ^ E5.5 embryonic epiblast and EpiLC nuclei compared to *Smcx*^fl/fl^ nuclei (Fig. 2a). These three genes were also transcribed from the Xist RNA-coated X-chromosome in significantly more *X*^ΔTsix^*Y* differentiating EpiLCs compared to *X*^ΔTsix^*X* females (Extended Data Fig. 2b-c). Together, the results in Figs. 1-2 demonstrate that the female-specific dose of SMCX is not only required for Xist RNA expression but also for efficient silencing of X-linked genes once Xist RNA is induced.

We next hypothesized that the *Smcx*^fl/Δ^ cells that displayed inefficient silencing express a reduced level of SMCX than the *Smcx*^fl/Δ^ cells with more stringent silencing of X-linked genes. Although *Smcx* escapes X-inactivation, expression of *Smcx* from the inactive-X is lower than from the active-X^21,22^ (Extended Data Fig. 2d). Therefore, *Smcx*^fl/Δ^ cells harboring the *Smcx*^fl^ allele on the inactive-X are expected to express reduced SMCX than those with the *Smcx*^fl^ allele on the active-X. If SMCX functions as a dose-dependent regulator of X-linked gene silencing, then the former group of cells would be more susceptible to X-inactivation defects than the latter (schematic in Fig. 2b).

To test this model, we derived hybrid WT *Smcx*^JF1/fl^ and heterozygous *Smcx*^JF1/Δ^ ESCs containing X-chromosomes from divergent strains of mice. In these cells, the X-chromosome harboring the *Smcx*^JF1^ allele (*X*^JF1^), which encodes a WT SMCX protein, is derived from the *Mus molossinus* JF1 strain, and the X-chromosome with the *Smcx*^fl^ or the *Smcx*^Δ^ allele (*X*^flSmcx^ and *X*^ΔSmcx^, respectively) is derived from the *Mus musculus* laboratory strain. We generated EpiLCs from these hybrid ESCs, and exploited single nucleotide polymorphisms (SNPs) in *Xist* on the two X-chromosomes to mark the X that was chosen for inactivation by allele-specific Xist RNA FISH (see Methods; Extended Data Fig. 3a). Xist RNA is upregulated from and coats the X-chromosome that is selected for inactivation^9,10,36^. We concurrently evaluated the expression of the three genes that were not efficiently silenced in *Smcx*^fl/Δ^ EpiLCs, *Lamp2*, *Mecp2*, and *Atrx*, via RNA FISH (see Fig. 2a). In the control *Smcx*^JF1/fl^ EpiLCs, the three genes were silenced with similar efficiency irrespective of which X-chromosome was Xist RNA coated (Fig. 2c), as expected, since both Xs harbor active *Smcx* alleles that generate WT protein. By contrast, in *Smcx*^fl/Δ^ EpiLCs all three genes were significantly more likely to be expressed from the Xist RNA-coated WT *X*^JF1^ as compared to the *X*^ΔSmcx^ (Fig. 2c; Extended Data Figs. 3b-f and 4). These data reiterate a dose-dependent function of SMCX in X-linked gene silencing during X-inactivation.

### Sufficiency of SMCX in Xist RNA induction and X-linked gene silencing

Having established a dose-dependent requirement of SMCX in X-linked gene silencing, we next tested the sufficiency of *Smcx* in inducing X-inactivation in male cells. Strikingly, ectopic expression of a mouse *Smcx* (*Tg*-*mSmcx*) as well as a human *SMCX* (*Tg-hSMCX*) transgene resulted in induction of Xist RNA in >40% of *XY* male ESCs (Fig. 3a). We additionally tested if the Xist RNA induction required the demethylase activity of SMCX. Expression of a point-mutant that abolishes the demethylase function of SMCX (*Tg-hSMCX(H514A)*)^15^ was not able to induce Xist RNA in the *XY* male ESCs (Fig. 3a). Similarly, ectopic expression of the closely related SMCX homolog, SMCY/KDM5D (*Tg*-*hSmcy*), which maps to the Y-chromosome and is itself capable of demethylating H3K4me2/3^15^, was also insufficient to activate Xist RNA expression (Fig. 3a). Thus, although both SMCX and SMCY can demethylate H3K4me2/3, only SMCX is able to induce *Xist* in an enzymatic activity-dependent manner (see Extended Data Fig. 5 and Supplemental Fig. 1). Moreover, more than 75% of the Xist RNA-coated *Tg*-*mSmcx* and *Tg-hSMCX XY* ESCs also silenced the three X-linked genes that are not efficiently silenced in *Smcx*^fl/Δ^ female EpiLCs, *Lamp2*, *Mecp2*, and *Atrx* (Figs. 3b and 2a), supporting the dual functions of SMCX in *Xist* induction and X-linked gene silencing.

We next took advantage of the sensitized background of *X*^ΔTsix^*Y* ESCs to further investigate the dose-dependency of SMCX in inducing *Xist* expression and X-linked gene silencing. In differentiating *X*^ΔTsix^*Y* male EpiLCs, some but not all cells ectopically induce Xist RNA due to a deletion in the *Xist* antisense repressor *Tsix*^14^; moreover, only a subset of the cells with Xist RNA coats silence X-linked genes (outlined in Fig. 3c). We first tested if SMCX overexpression could increase the frequency of Xist RNA-coated *X*^ΔTsix^*Y* nuclei. Indeed, compared to the parent *X*^ΔTsix^*Y* cells, d2 differentiated EpiLCs ectopically expressing mouse and human SMCX (*Tg-mSmcx*;*X*^ΔTsix^*Y* and *Tg-hSmcx*;*X*^ΔTsix^*Y*, respectively) displayed a significant increase in the percentage of Xist RNA-coated nuclei (Fig. 3d-e and Extended Data Fig. 6a). Cells ectopically expressing the enzymatically-inactive SMCX protein (*Tg-hSMCX(H514A)*;*X*^ΔTsix^*Y*), SMCY (*Tg-hSMCY*;*X*^ΔTsix^*Y*), or lacking the Y-chromosome (*X*^ΔTsix^*O*), on the other hand, exhibited a similar frequency of Xist RNA-coated X-chromosomes as the *X*^ΔTsix^*Y* cells (Fig. 3d). Reciprocally, the loss of SMCX in *X*^ΔTsix;ΔSmcx^*Y* and *X*^ΔTsix;ΔSmcx^*O* cells reduced the incidence of ectopic Xist RNA coating (Fig. 3d). Xist RNA levels quantified by RT-qPCR were concordant with the RNA FISH data (Fig. 3e).

In addition to Xist RNA induction, we examined the impact of SMCX in the silencing of X-linked genes in *X*^ΔTsix^*Y* cells. The overexpression of SMCX in differentiated *X*^ΔTsix^*Y* EpiLCs resulted in an increased frequency of X-linked gene silencing (Fig. 3f; Extended Data Fig. 6b-c and Extended Data Table 1). Increased expression of the mutant SMCX (H514A) or SMCY again resulted in a gene silencing pattern similar to the parental *X*^ΔTsix^*Y* cells. Conversely, the absence of SMCX in differentiating *X*^ΔTsix;ΔSmcx^*Y* and *X*^ΔTsix;ΔSmcx^*O* EpiLCs significantly reduced the percentage of nuclei with silenced X-linked genes. Moreover, *X*^ΔTsix^*O* and *X*^ΔTsix;ΔSmcx^*O* differentiating EpiLCs recapitulated the pattern of silencing in differentiating *X*^ΔTsix^*Y* and *X*^ΔTsix;ΔSmcx^*Y* cells, thus excluding a contribution of Y-linked genes in the observed differences.

### SMCX dose-dependent histone H3K4 demethylation on the inactive-X

To test if SMCX directly regulates X-inactivation, we first examined if SMCX is enriched on the inactive-X cytologically. To ensure efficient detection of SMCX, we generated *XX* female ESCs stably expressing an epitope-tagged SMCX protein (Strep-SMCX). Strep-SMCX accumulated on the X-chromosome as Xist RNA began to coat the X-chromosome prior to the differentiation of ESCs into EpiLCs (Fig. 4a). SMCX enrichment was transient and was largely lost by the time the ESCs differentiated into EpiLCs. In addition, only the Xist RNA-coated inactive X-chromosome, but not the active-X, displayed the SMCX accumulation in the female cells (Fig. 4a), indicating that the transient SMCX recruitment is an Xist RNA-dependent process. A similarly transient accumulation of SMCX characterized the onset of Xist RNA accumulation in differentiating *X*^ΔTsix^*Y* EpiLCs. Slower kinetics of the SMCX enrichment on the X-chromosome upon Xist RNA coating in male compared to female cells (Fig. 4a) mirrors the slower induction of Xist RNA in the *X*^ΔTsix^*Y* cells^37^.

SMCX is a member of the KDM5-family of enzymes that demethylate H3K4me2/3^15,38^, which are well-characterized hallmarks of transcriptionally-engaged chromatin^17^. H3K4me2 is excluded from the inactive-X^20^. To test if SMCX is responsible for H3K4me2/3 exclusion, we examined H3K4me2 depletion from the inactive-X as a function of SMCX dose. The percentage of nuclei exhibiting depletion of H3K4me2 and H3K4me3 from the Xist RNA-coated X-chromosome increased with increasing SMCX dose (Fig. 4b and Extended Data Fig. 7a). To further probe H3K4me2 occupancy on the inactive-X when SMCX dose is modulated, we performed H3K4me2 ChIP-Seq in differentiating *X*^ΔTsix^*Y*, *X*^ΔTsix;ΔSmcx^*Y*, and *Tg-Smcx*;*X*^ΔTsix^ EpiLCs. We found an inverse correlation between cellular SMCX levels and H3K4me2 genome-wide (Fig. 4c-d). The increase in H3K4me2 upon the loss of SMCX was more pronounced on the X-chromosome compared to the autosomes, suggesting that H3K4me2 occupancy on the X-chromosome is more sensitive to a reduced SMCX dose compared to autosomes (Fig. 4c and Extended Data Fig. 7b-d). To investigate the relationship of H3K4me2 and transcription, we examined H3K4me2 at TSSs of the eight X-linked genes profiled by RNA FISH in Fig. 2a. The three genes that were not silenced stringently in female *Smcx*^fl/Δ^ cells, *Lamp2*, *Mecp2*, and *Atrx*, showed the greatest change in promoter H3K4me2 levels as a function of SMCX dose (Fig. 4e and Extended Data Fig. 8a). Thus, X-linked gene silencing by SMCX entails the removal of H3K4me2 from promoters.

The impact of SMCX dose on H3K4me2 levels persisted even after the transient SMCX enrichment on the X-chromosome dissipated (Fig. 4a-e). Although the cytological accumulation of SMCX on the inactive-X disappeared in differentiating EpiLCs (Fig. 4a), by ChIP-Seq residual SMCX protein was nevertheless detected at gene regulatory elements such as promoters and distal DNaseI hypersensitive sites (DHS) at X-linked genes (Extended Data Fig. 8a-b). At the SMCX-enriched promoters, loss of SMCX led to a significant increase in H3K4me2 at both the X-chromosome as well as on a genome-wide scale (Extended Data Fig. 8c). These results suggest that SMCX is transiently recruited onto the X-chromosome at the onset of X-inactivation and that the SMCX-dependent removal of H3K4me2 at promoters is maintained after SMCX dislodges from X-linked genes.

We then sought to probe the mechanism by which SMCX induces *Xist* expression. In contrast to the TSS-surrounding SMCX ChIP-Seq signals found at most X-linked genes, at the *Xist* locus, SMCX is enriched ∼1.5 kb downstream of the TSS (Fig. 5a-b and Extended Data Fig. 8a). We previously reported an enhancer-like regulatory element that promotes *Xist* expression in this region^39^. Importantly, SMCX reverses H3K4me2/3 and leaves intact H3K4me1^15^, which is a hallmark of transcriptional enhancers^40,41^. Active enhancers are also enriched in H3K27 acetylation (H3K27ac)^42^. We therefore tested H3K4me1/2/3 and H3K27ac occupancy at the putative *Xist* enhancer by ChIP-qPCR.

Consistent with the previous study^39^, we noted that the SMCX-enriched region downstream of the *Xist* TSS indeed displayed enhancer-like chromatin signature of high H3K4me1 and lower H3K4me2/3 in d2 differentiated *X*^ΔTsix^*Y* EpiLCs (Fig. 5c). Conversely, the TSS-proximal region of *Xist* was decorated with higher H3K4me2/3 and lower H3K4me1. In the *Xist* enhancer region, SMCX overexpression led to a further increase in H3K4me1 and a decrease in H3K4me2/3, and was accompanied by appearance of the active enhancer mark H3K27ac. In contrast, at the *Xist* promoter, which is devoid of SMCX, we observed an increase in the active chromatin signatures H3K4me2/3 and H3K27ac upon SMCX overexpression. When we analyzed the *Atrx* promoter, which is directly bound by SMCX in the d2 differentiating EpiLCs, SMCX overexpression resulted in the removal of H3K4me2/3. These changes in histone modification levels at the *Xist* enhancer or the *Atrx* promoter were not observed when we overexpressed the mutant SMCX-H514A mutant (Fig. 5c), consistent with the demethylase activity-dependent induction of Xist RNAs (Fig. 4). The absence of SMCX resulted in an opposite pattern in histone modifications compared to SMCX overexpression, reinforcing the SMCX-dose dependency in both promoter and enhancer regulation (Fig. 5c). Note that *Tsix* is not expressed in the *X*^ΔTsix^*Y* EpiLCs^33^; therefore, we rule out indirect induction of SMCX via *Tsix* suppression. These results support a model in which SMCX directly induces *Xist* expression via enhancer activation and silences X-linked genes by promoter suppression.

## Discussion

In this study, we show that the histone H3K4me2/3 demethylase SMCX functions in a dose- and enzymatic activity-dependent manner both to induce *Xist* and, separately, to silence X-linked genes (Fig. 5d). The higher, biallelic expression of SMCX links the discrete steps that are thought to underlie random X-inactivation^43,44^: sensing the X-chromosome complement in the cell (‘counting’ step), inducing expression of *Xist* (‘initiation’ step), and silencing of X-linked genes (‘establishment’ step). All X-chromosome sequence elements necessary and sufficient to induce X-inactivation are thought to reside within the *X-inactivation center* (*Xic*), which harbors the *Xist* locus^43,44^. *Smcx*, however, maps far outside of the *Xic*, near the telomeric end of the X-chromosome. The results herein argue against the primacy of the *Xic* and instead position *Smcx* at or near the top of the molecular hierarchy leading to random X-inactivation (Fig. 5d).

X-inactivation is postulated to have evolved as a consequence of the deterioration of the Y-chromosome^2^, which was once homologous to the X-chromosome in the common ancestor of therian mammals^45^. By extension, the presence of active Y-linked genes is believed to have driven escape from inactivation of the X-linked homologs to adjust the dose of X-Y gene pairs between the sexes^46^. Our data instead demonstrate that X-Y homologs can be functionally distinct, with the X-linked copy having evolved a female-specific function to cause X-inactivation. Escape from X-inactivation may therefore be evolutionarily driven in part by the requirement of the X-linked homologs in X-chromosome dosage compensation.

**Figure 1.**
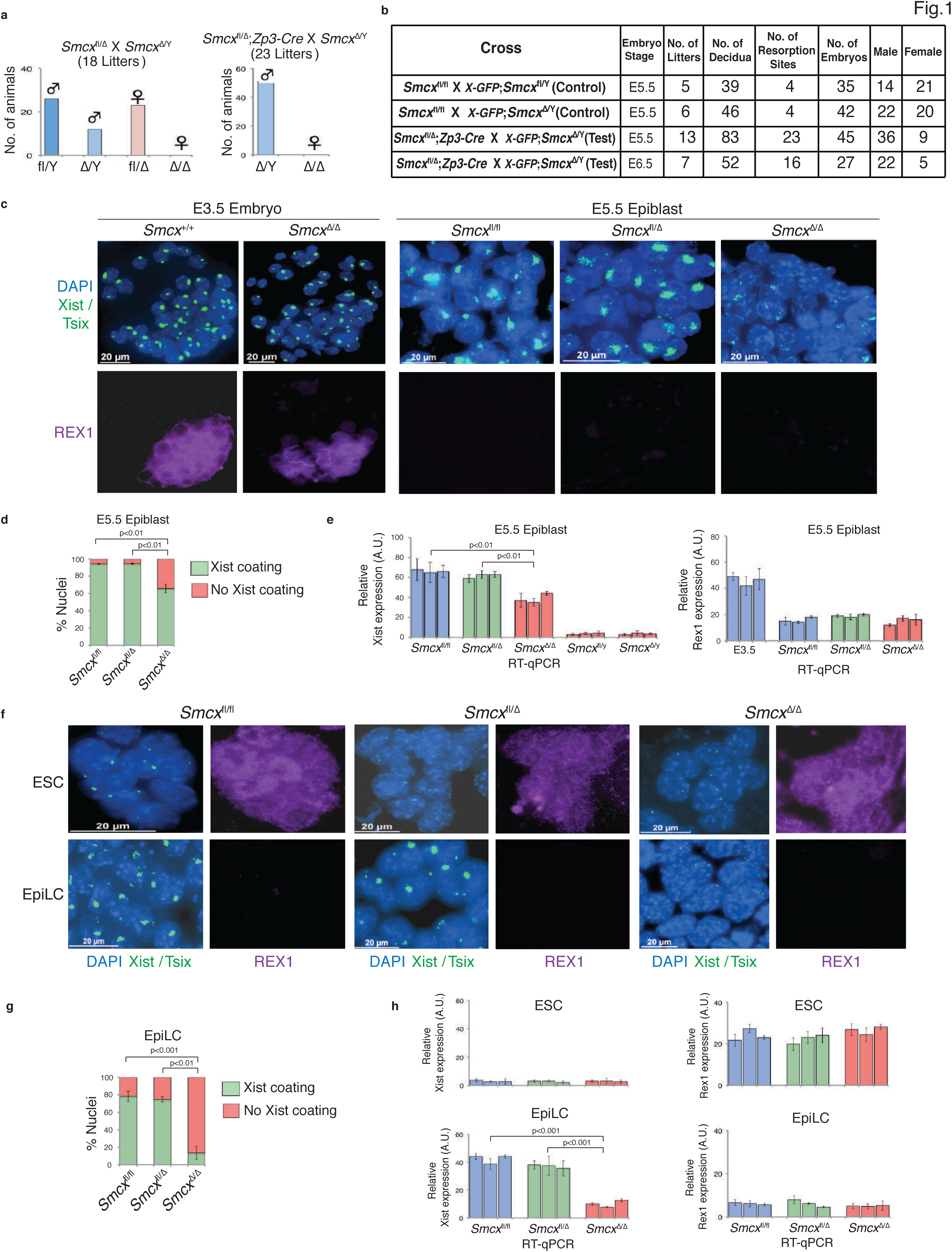
Defective X-inactivation in *Smcx*^Δ/Δ^ female embryos and EpiLCs. **a-b,** Live-born animals and E5.5 and E6.5 embryos obtained from intercrosses of *Smcx*-mutant animals. **c,** RNA FISH detection of Xist/Tsix RNAs coupled with IF staining for the pluripotency marker REX1 in representative E5.5 epiblasts. *Rex1* downregulation marks differentiating pluripotent epiblast progenitors^34,47^. Nuclei are stained blue with DAPI. **d**, Quantification of Xist RNA coated nuclei in the E5.5 epiblasts. n=150-315 nuclei in each of 3 embryos/genotype. **e,** Relative quantification of Xist and Rex1 RNAs by RT-qPCR in E5.5 epiblasts. n=3 embryos/genotype, each analyzed in triplicate. **f,** Xist/Tsix RNAs and REX1 profiling in representative ESCs and EpiLCs. **g,** Quantification of Xist RNA-coated EpiLCs. n=100 nuclei from EpiLCs differentiated from each of 3 independent ESC lines/genotype in **f-g**. **h,** Relative quantification of Xist and Rex1 RNAs by RT-qPCR in the ESCs and EpiLCs. n=3 independent ESC lines differentiated into EpiLCs/genotype, each analyzed in triplicate. Error bars, standard error. *p*-values, Welch’s two-sided t-test.

**Figure 2.**
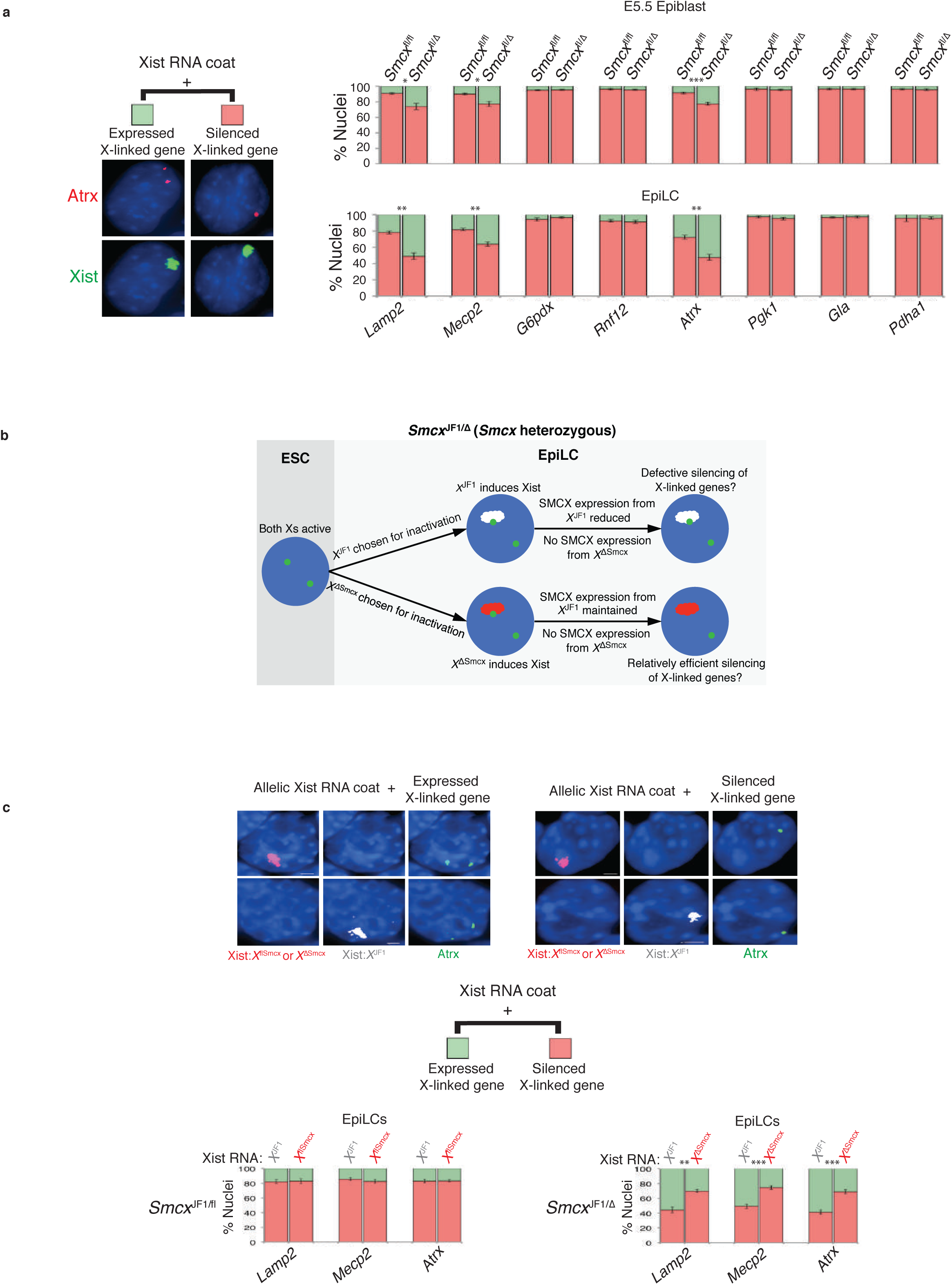
SMCX dose-dependent X-linked gene silencing in *Smcx*^fl/Δ^ EpiLCs. **a,** RNA FISH quantification of X-linked gene expression. Left, a representative nucleus with nascent transcripts of one of the genes surveyed, *Atrx*, in red and Xist RNA in green. n=86-141 nuclei in each of 3 embryonic epiblasts/genotype; and, n=100 nuclei from EpiLCs differentiated from each of 3 independent ESC lines/genotype. Error bars, standard error. *, *p* ≤ 0.05; \*\**, p* < 0.01; ***, *p* < 0.001, Welch’s two-sided t-tests. **b,** Schematic of possible patterns of X-linked gene expression by RNA FISH in *Smcx*^JF1/Δ^ EpiLCs. **c,** Top, representative example of the RNA FISH outcome in *Smcx*^JF1/Δ^ EpiLCs. Bottom, quantification of expression of three X-linked genes coincident with allele-specific Xist RNA coating in *Smcx*^JF1/fl^ or *Smcx*^JF1/Δ^ EpiLCs. n=100 nuclei from EpiLCs differentiated from each of 3 independent ESC lines/genotype.

**Figure 3.**
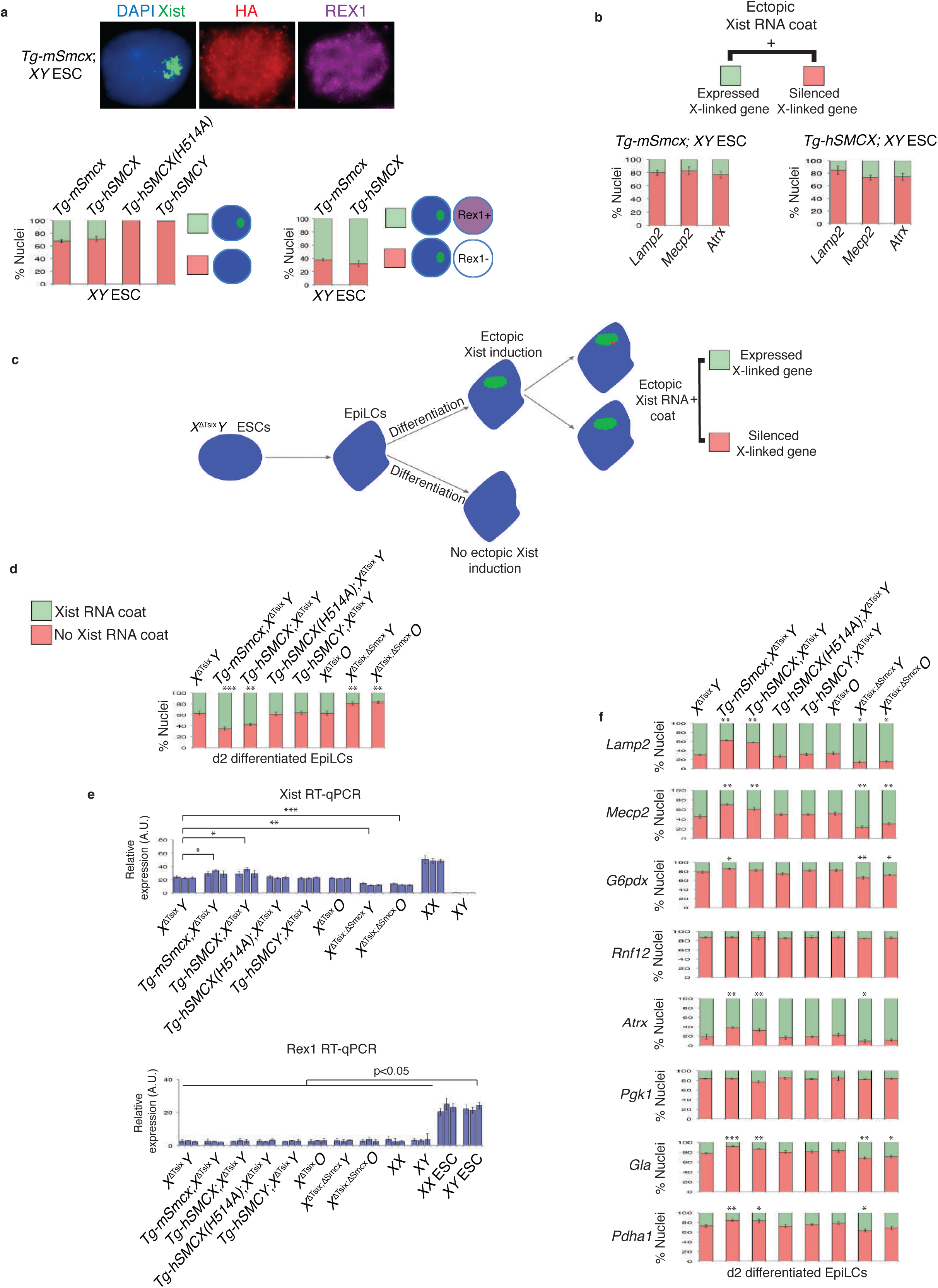
Ectopic Xist RNA induction upon SMCX overexpression in male cells. **a,** Representative example (top) and quantification (bottom) of RNA FISH detection of Xist/Tsix RNAs (green) coupled with IF staining for the HA epitope tag (red) and REX1 (purple) in *XY* male ESCs expressing an inducible mouse HA-SMCX transgene (*Tg-mSmcx*); human HA-SMCX transgene (*Tg-hSMCX*); an HA-SMCX mutant transgene lacking demethylase activity (*Tg-hSMCX(H514A)*); and, an HA-SMCY transgene (*Tg-hSMCY*). n=100 nuclei from each of 3 independent ESC lines. **b,** Expression of the X-linked genes *Lamp2*, *Mecp2*, and *Atrx* from the Xist RNA-coated X-chromosome in the transgenic *XY* ESCs. n=100 nuclei from each of 3 independent ESC lines. **c,** Schematic of ectopic Xist RNA coating and X-linked gene silencing during differentiation of *X*^ΔTsix^*Y* ESCs. **d,** Quantification of ectopic Xist RNA-coated nuclei in d2 differentiated EpiLCs. n=100 nuclei from EpiLCs differentiated from each of 3 independent ESC lines/genotype. **e,** Relative quantification of Xist (top) and Rex1 (bottom) RNAs by RT-qPCR in the d2 differentiated EpiLCs. n=3 independent ESC lines differentiated into EpiLCs/genotype, each analyzed in triplicate. **f,** Quantification of X-linked gene expression upon ectopic *Xist* induction in d2 differentiated EpiLCs. n=100 nuclei from EpiLCs differentiated from each of 3 independent ESC lines/genotype. Error bars, standard error. *, *p* ≤ 0.05; \*\**, p* < 0.01; ***, *p* < 0.001, Welch’s t-tests. All samples statistically compared to the percentage in *X*^ΔTsix^*Y* cells (see Extended Data Table 1 for exact values).

**Figure 4.**
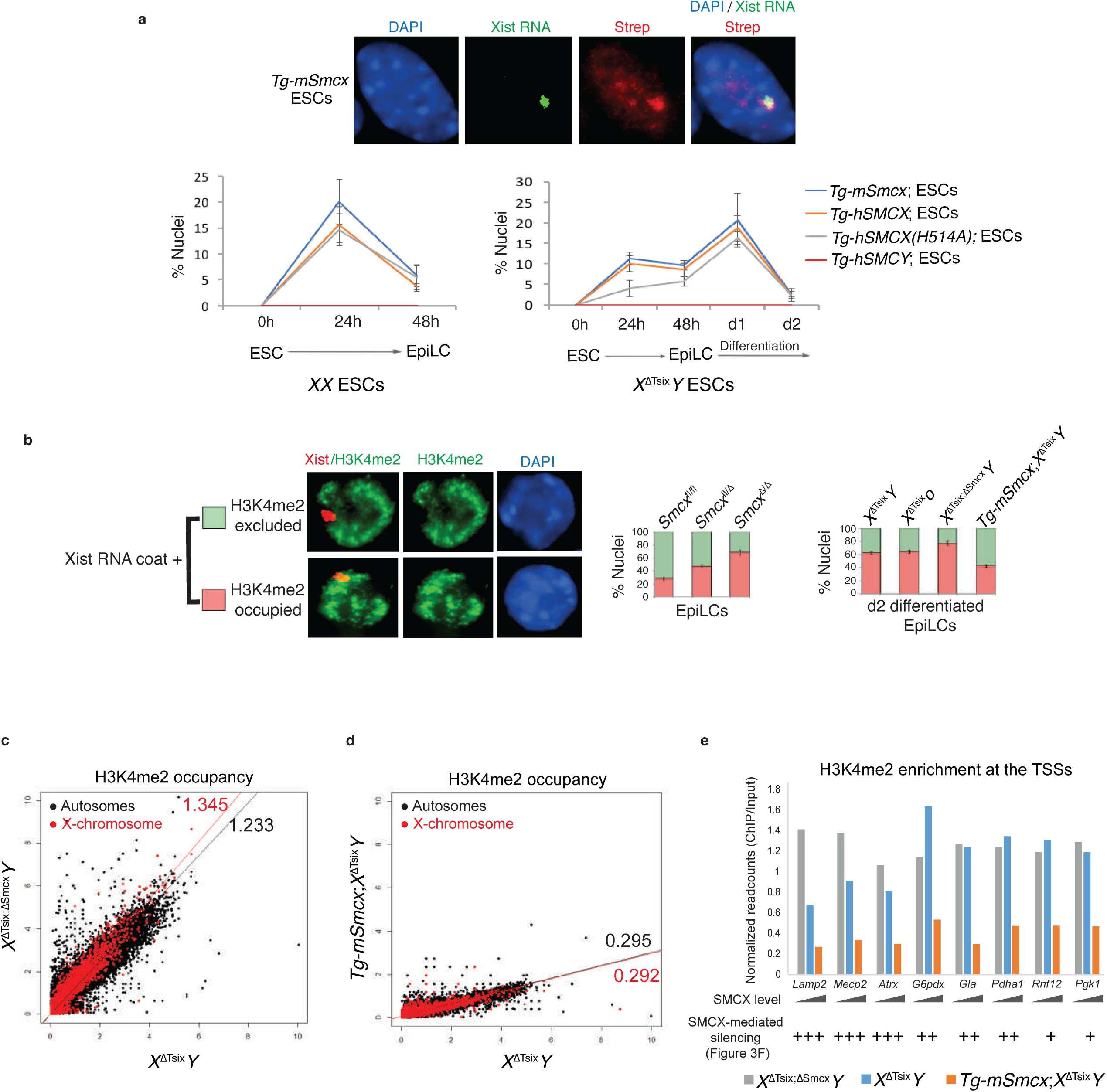
SMCX-dose dependent H3K4me2 exclusion from the inactive X-chromosome. **a,** SMCX is transiently enriched on the inactive-X. Representative example (top) and quantification (bottom) of RNA FISH detection of Xist RNA (green) coupled with detection the Strep-SMCX (red) in *XX* and *X*^ΔTsix^*Y* ESCs expressing mouse SMCX transgene (*Tg-mSmcx*); human Strep-SMCX transgene (*Tg-hSMCX*); Strep-SMCX mutant transgene lacking demethylase activity (*Tg-hSMCX(H514A)*); and, Strep-SMCY transgene (*Tg-hSMCY*). n=100 nuclei from each of 3 independent ESC lines. **b,** H3K4me2 exclusion from the Xist RNA-coated X-chromosome. Left, schematic and observed micrographs. Right, quantification of the data. **c-d,** H3K4me2 occupancy by ChIP-Seq with varying SMCX dose. Normalized H3K4me2 ChIP signals plotted over every 1 kb bin of the mouse genome (see Methods and Extended Data Fig. 7b-f). **e,** Normalized H3K4me2 ChIP-Seq read counts ±1 kb of transcription start sites (TSSs) for the eight X-linked genes profiled by RNA FISH in Figs. 2a and 3f. The SMCX-dose dependent H3K4me2 reduction was more evident in *Lamp2*, *Mecp2*, and *Atrx*, whose expression is also more sensitive to SMCX levels (Figs. 2a and 3f). Plots in **c-e** are from two independent differentiating EpiLC lines.

**Figure 5.**
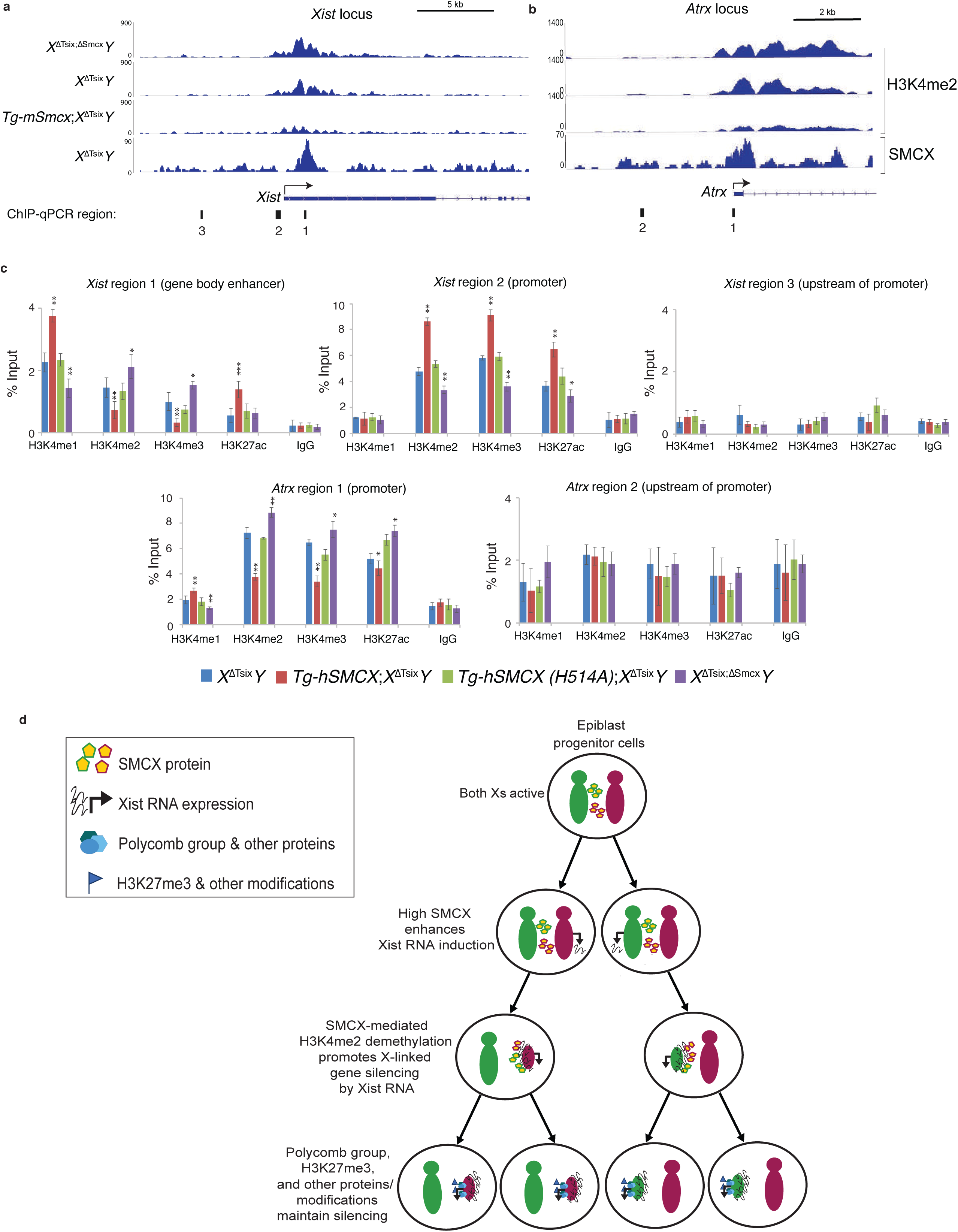
SMCX induces *Xist* expression via enhancer activation. **a,** SMCX and H3K4me2 occupancy at the *Xist* locus. H3K4me2 enrichment downstream of the *Xist* TSS is inversely correlated to SMCX dose and *Xist* expression. **b,** SMCX and H3K4me2 occupancy at the *Atrx* locus. **c,** ChIP-qPCR quantification of H3K4me1, H3K4me2, H3K4me3, and H3K4ac occupancy at the *Xist* and *Atrx* loci in d2 differentiated EpiLCs. Locations of PCR amplicons are denoted beneath the browser views in **a** and **b**. Mean ± SEM. *, *p* ≤ 0.05; \*\**, p* < 0.01; ***, *p* < 0.001, Welch’s t-tests. **d,** A model of SMCX function in X-inactivation.

## Methods

### Ethics Statement

This study was performed in strict accordance with the recommendations in the Guide for the Care and Use of Laboratory Animals of the National Institutes of Health. All animals were handled according to the protocols approved by the University Committee on Use and Care of Animals (UCUCA) at the University of Michigan (protocol #PRO00004007).

### Mice

The generation of *Smcx*^fl/fl^ mice and mice harboring the *Tsix*^AA2Δ1.7^ (*Tsix*^Δ^) mutation have been described previously^25,48^. The X-linked *GFP* transgenic (*X-GFP*) and JF1 strains have also been described previously^30,31,49,50^.

### Embryo Dissections and Processing

Pre-, peri-, and post-implantation stage embryos were isolated essentially as described previously^51^. Briefly, E3.5 embryos were flushed from the uterine limbs in 1X PBS (Invitrogen, #14200075) containing 6 mg/ml bovine serum albumin (BSA; Invitrogen, #15260037). The zona pellucidas surrounding E3.5 embryos were removed through incubation in cold Acidic Tyrode’s Solution (Sigma, #T1788), followed by neutralization through several transfers of cold M2 medium (Sigma, #M7167). GFP fluorescence conferred by the paternal transmission of the *X-GFP* transgene was used to distinguish female from male embryos, since only females inherit the paternal X-chromosome. Isolated embryos were either lysed for RNA isolation or plated onto gelatin-coated glass coverslips in 1X PBS with 6 mg/ml BSA for immunofluorescence (IF)/RNA fluorescence *in situ* hybridization (FISH) staining. For plated embryos, excess solution was aspirated, and coverslips were air-dried for 15 min. After drying, embryos were permeabilized and fixed in 50 μL solution of 0.05% Tergitol (Sigma, #NP407) and 1% paraformaldehyde (Electron Microscopy Sciences, #15710) in 1X PBS for 10 min. Excess solution was tapped off, and coverslips were rinsed 3X with 70% ethanol and stored in 70% ethanol at −20°C prior to RNA FISH. For isolation of post-implantation embryos (E5.5, E6.5 and E7.5), individual implantation sites were cut from the uterine limbs and decidua were removed with forceps in 1X PBS/6 mg/ml BSA. Embryos were dissected from the decidua, and the Reichert’s membranes surrounding post-implantation embryos were removed using fine forceps. For separation of extra-embryonic and epiblast portions of embryos, fine forceps were used to physically bisect the embryos at the junction of the epiblast and extra-embryonic ectoderm. The epiblast was further distinguished by GFP fluorescence conferred by the paternally-transmitted *X-GFP* transgene; the transgene is mosaically expressed in the epiblast due to random X-inactivation but is silenced in the extra-embryonic tissues because of imprinted X-inactivation of the paternal-X^30,31,49,50^. Extra-embryonic and embryonic epiblast cells were then separately either lysed for RNA isolation or plated in 0.25X PBS with 6 mg/mL BSA onto gelatinized coverslips for IF/RNA FISH staining. Samples plated for IF/RNA FISH were fixed, permeabilized, and stored as described for E3.5 embryos.

### Derivation, Culture and Differentiation of Embryonic Stem Cell (ESCs) Lines

ESC lines were derived following the protocol previously described^52^. Cells were cultured on mouse embryo fibroblasts (MEFs) in Knockout DMEM (GIBCO, #10829-018) with 15% Knockout Serum Replacement (GIBCO, #A1099201), 5% FBS (GIBCO, #104390924), 2 mM L-glutamine (GIBCO, #25030), 0.1 mM 2-mercaptoethanol (Sigma, #M7522), 1X nonessential amino acids (GIBCO, #11140-050), supplemented with 1 μM GSK3 inhibitor CHIR99021 (Stemgent #04-0004), 1 μM MEK inhibitor PD0325901 (Stemgent #04-0006) and 1000 units/mL LIF (Millipore #ESG1106). The *X*^ΔTsix^*O* ESC lines were subcloned from *X*^ΔTsix^*X* ESC lines. The *X*^ΔTsixΔSmcx^*O* ESC lines were subcloned from *X*^ΔTsixΔSmcx^*X* ESC lines. The *X*^ΔSmcx^*O* and *X*^JF1^*O* ESC lines were subcloned from *X*^JF1^*X*^ΔSmcx^ ESC line. The loss of the second X-chromosome in the *X*^ΔSmcx^*O* and *X*^JF1^*O* ESC lines was ascertained via RNA and DNA FISH for *Xist* and the eight X-linked genes depicted in Fig. 2a.

For IF and/or RNA FISH, cells were then permeabilized through sequential treatment with ice-cold cytoskeletal extraction buffer (CSK:100 mM NaCl, 300 mM sucrose, 3 mM MgCl2, and 10 mM PIPES buffer, pH 6. 8) for 30 sec, ice-cold CSK buffer containing 0.4% Triton X-100 (Fisher Scientific, #EP151) for 30 sec, followed twice with ice-cold CSK for 30 sec each. After permeabilization, cells were fixed by incubation in 4% paraformaldehyde for 10 min. Cells were then rinsed 3X in 70% ethanol and stored in 70% ethanol at −20°C prior to IF and/or RNA FISH.

### Differentiation of ESCs into Epiblast-like Cells (EpiLCs)

To convert ESC to EpiLC, ESC lines were grown in 2i culture conditions (N2B27 medium consisting 50% DMEM/F12, 50% neurobasal media, 2 mM L-glutamine (GIBCO, #25030), 0.1 mM 2-mercaptoethanol (Sigma, #M7522), N2 supplement (Invitrogen #17502048), B27 supplement (Invitrogen #17504-044), supplemented with 1 μM GSK3 inhibitor CHIR99021 (Stemgent #04-0004), 1 μM MEK inhibitor PD0325901 (Stemgent #04-0006), and 1000 U/ml LIF (Millipore #ESG1106) in gelatin-coated tissue culture dishes for 4 passages (Buecker et al., 2014; Hayashi et al., 2011). To achieve EpiLC differentiation, cells were cultured in N2B27 medium supplemented with 10 ng/ml FGF2 (R&D Systems, #233-FB) and 20 ng/ml Activin A (R&D Systems, #338-AC) in Fibronectin (15μg/ml) (Sigma #F1141)-coated tissue culture dishes for 48 hrs. For further differentiation, the cells were cultured in N2B27 medium without FGF2 and Activin A for an additional 2, 4 and 6 days (d2, d4 and d6 of differentiation, respectively). For IF and/or RNA FISH, cells were permeabilized, fixed, and CSK processed as described above for ESCs.

### Derivation and Culture of Epiblast Stem Cell (EpiSC) Lines

EpiSCs (*X*^ΔTsix^*X*^JF1^ #2, #15) were derived and characterized in a previous study^33^. For derivation of the EpiSCs lines, the epiblast layer was microdissected from E5.5 embryos and plated on MEF cells in EpiSC medium consisting of Knockout DMEM (GIBCO, #10829-018) supplemented with 20% Knockout Serum Replacement (KSR; GIBCO, #A1099201), 2 mM L-glutamine (GIBCO, #25030), 1X nonessential amino acids (GIBCO, #11140-050), and 0.1 mM 2-mercaptoethanol (Sigma, #M7522), 10 ng/ml FGF2 (R&D Systems, #233-FB) and cultured for 3-4 days to form a large EpiSC colony. EpiSC colonies were then manually dissociated into small clusters using a glass needle and plated into 1.9 cm^2^ wells containing MEF feeders in EpiSC cell medium. EpiSCs were passaged every third day using 1.5 mg/ml collagenase type IV (GIBCO, #17104-019) with pipetting into small clumps.

### RNA Fluorescence *in situ* Hybridization (RNA FISH)

RNA FISH with double-stranded and strand-specific probes was performed as previously described^14,49,53^. The dsRNA FISH probes were made by randomly-priming DNA templates using BioPrime DNA Labeling System (Invitrogen, #18094011). Strand-specific *Xist* probes were generated from templates as described^39,51^. Probes were labeled with Fluorescein-12-dUTP or -UTP (Invitrogen) or Cy3-dCTP or –CTP (GE Healthcare). Labeled probes from multiple templates were precipitated in a 0.3M sodium acetate solution (Teknova, #S0298) along with 300 μg of yeast tRNA (Invitrogen, #15401-029) and 150 μg of sheared, boiled salmon sperm DNA (Invitrogen, #15632-011). The solution was then spun at 15,000 rpm for 20 min at 4°C. The pellet was washed consecutively with 70% ethanol and 100% ethanol. The pellet was dried and re-suspended in deionized formamide (ISC Bioexpress, #0606). The probe was denatured by incubating at 90°C for 10 min followed by an immediate 5 min incubation on ice. A 2X hybridization solution consisting of 4X SSC, 20% Dextran sulfate (Millipore, #S4030), and 2.5 mg/ml purified BSA (New England Biolabs, #B9001S) was added to the denatured solution. All probes were stored at −20°C until use.

Cells or embryo fragments mounted on coverslips were dehydrated through 2 min incubations in 70%, 85%, 95%, and 100% ethanol solutions and subsequently air-dried. The coverslips were then hybridized to the probe overnight in a humid chamber at 37°C. The samples were then washed 3X for 7 min each while shaking at 39°C with 2XSSC/50% formamide, 2X with 2X SSC, and 2X with 1X SSC. A 1:250,000 dilution of DAPI (Invitrogen, #D21490) was added to the third 2X SSC wash. Coverslips were then mounted on slides in Vectashield (Vector Labs, #H-1200).

### Allele-specific Xist RNA FISH

Allele specific Xist RNA FISH were generated as described by Levesque *et al.* (2013)^54^. Briefly, a panel of short oligonucleotide probes were designed to uniquely detect either the *M. musculus* or the *M. molossinus* alleles of *Xist*. Five probes were designed for each *Xist* allele; each probe overlaps a SNP that differs between the two strains, with the SNP located at the fifth base pair position from the 5’ end. The same panel of five SNPs was used for both sets of allele-specific probes. The 3’ end of each oligonucleotide probe is fluorescently tagged using Quasar dyes (Biosearch technologies): *M. musculus*-specific oligos were labeled with Quasar 570 and *M. molossinus* oligos labeled with Quasar 670. In addition to labeled SNP-overlapping oligonucleotides, a panel of 5 “mask” oligonucleotides was also synthesized. These “mask” probes are complimentary to the 3’ end of the labeled allele-specific probes and will initially hybridize to the allele-specific oligonucleotides, leaving only 9-10 base pairs of sequence surrounding the polymorphic site available to initially hybridize to the target Xist RNA; since this region is short, the presence of a single nucleotide polymorphism is sufficient to destabilize unintended hybridization with the alternate allele. Sequences of detection and mask probes are listed in Methods Table 1 below. The allele-specific Xist RNA FISH probes were tested together with a strand-specific Xist RNA probe, labeled with Fluorescein-12-UTP (Invitrogen), which served as a guide probe that hybridized to Xist RNA generated from both *Xist* alleles and ensured that the allele-specific probes were faithfully detecting Xist RNA. The guide Xist RNA probe was first ethanol precipitated as previously described, then resuspended in hybridization buffer containing 10% dextran sulfate, 2X saline-sodium citrate (SSC) and 10% formamide. The precipitated guide RNA probe was then mixed with the *M. musculus* and *M. molossinus* detection probes, to a final concentration of 5 nM per allele-specific oligo, and 10 nM mask probe, yielding to 1:1 mask:detection oligonucleotide ratios. Coverslips were hybridized to the combined probe overnight in a humid chamber at 37°C. After overnight hybridization, samples were washed twice in 2X SSC with 10% formamide at 37°C for 30 min, followed by two washes in 2X SSC for 5 mins. A 1:250,000 dilution of DAPI (Invitrogen, #D21490) was added to the second 2X SSC wash. Coverslips were then mounted on slides in Vectashield (Vector Labs, #H-1200).

### DNA FISH

DNA FISH probes were prepared as described for double-stranded RNA-FISH probes^14,53^. The BAC template used for *Xist* DNA FISH is RP24-287F13 (Children’s Hospital of Oakland Research Institute). After RNA FISH, cells were washed with 1X PBS three times and then incubated in PBS for 5 min at room temperature. The cells were then refixed with 1% (wt/vol) PFA containing 0.5% (vol/vol) Tergitol and 0.5% (vol/vol) Triton X-100 for 10 min at room temperature. The cells were next dehydrated through an ethanol series (70%, 85%, and 100% ethanol, 2 min each) and air dried for 15 mins. The cells were then treated with RNase A (1.25 μg/μl) at 37°C for 30 min. The cells were again dehydrated through the ethanol series as described above. The samples were then denatured in a prewarmed solution of 70% formamide in 2X SSC on a glass slide stationed on top of a heat block set at 95°C for 11 min followed immediately by dehydration through a −20°C-chilled ethanol series (70%, 85%, 95%, and 100% ethanol, 2 min each). The cells were then air dried for 15 min followed by probe hybridization overnight at 37°C. The following day, the samples were washed twice with prewarmed 50% formamide/2X SSC solution at 39°C and 2X with 2X SSC, 7 min each.

### Immunofluorescence (IF)

Cells mounted on glass coverslips were washed 3X in PBS for 3 min each while shaking. Coverslips were then incubated in blocking buffer consisting of 0.5 mg/mL BSA (New England Biolabs, #B9001S), 50 μg/mL yeast tRNA (Invitrogen, #15401-029), 80 units/mL RNAseOUT (Invitrogen, #10777-019), and 0.2% Tween 20 (Fisher, #BP337-100) in 1X PBS in a humid chamber for 30 min at 37°C. The samples were next incubated with primary antibody diluted in blocking buffer for 1-3 hr in the humid chamber at 37°C. Anti-REX1 antibody (Thermo Scientific, #PA5-27567) was used at 1:150 dilution; anti-HA antibody was used at 1:100 dilution (Abcam, #18181); anti-H3K4me2 was used at 1:200 (Millipore, #07030); and, the anti-H3S10ph antibody (Cell Signaling, #9701) was used at 1:150 dilution. The Cleaved Caspase-3 (ASP175) (5A1E) antibody (Cell Signaling # 9664) was used at 1:200. The samples were then washed 3X in PBS/0.2% Tween 20 for 3 min each while shaking. After a 5 min incubation in blocking buffer at 37°C in the humid chamber, the samples were incubated in blocking buffer containing a 1:300 dilution of fluorescently-conjugated secondary antibody (Alexa Fluor, Invitrogen) for 30 min in the humid chamber at 37°C, followed by three washes in PBS/0.2% Tween 20 while shaking for 3 min each. The samples were then processed for RNA FISH.

### RT-qPCR

RT-qPCR was performed using SYBR Green-based relative quantification method on an Eppendorf Realplex Mastercycler. The housekeeping gene *Gapdh* was used as an internal control for data normalization. cDNA was synthesized using SuperScript III First Strand Synthesis System (Invitrogen # 18080-051), following manufacturer’s instructions. Control reactions lacking reverse transcriptase for each sample were also performed to rule out genomic DNA contamination. Real time PCR was carried out using the SYBR Green PCR Mastermix (Kapa Biosystem #KK4650). The following primers were used for RT and PCR: *Smcx*: forward TTTGGCAGCGGTTTCCCTGTCAGT, reverse AAGACCATTCCCACATACAGCC; *Rex1*: forward CCACAAACAGATCCGGCTT, reverse TGGAAGCGAGTTCCCTTCTC; *Xist*: forward CAAGAAGAAGGATTGCCTGGATTT, reverse GCGAGGACTTGAAGAGAAGTTCTG. *P*-values for all RT-qPCR results were calculated using Welch’s two-sample *T*-tests.

### Quantification of Allele-specific Expression by Pyrosequencing

Allele-specific expression of *Smcx* was quantified using Qiagen PyroMark sequencing platform. An *Smcx* RT-PCR amplicon containing a single nucleotide polymorphism (SNP) was designed using PyroMark Assay Design software. Single cell lysates were prepared in 10 μL of lysis buffer from the Single Cell RT-PCR Assay Kit by Signosis (# CL-0002). DNA was removed from samples by spinning at 12000 rpm for 5 min, followed by DNase I treatment (37°C for 30 min). Five μl of cell lysate was then used directly for cDNA synthesis with the Invitrogen SuperScript III One-Step RT-PCR System (#12574-026). Control reactions lacking reverse transcriptase for each sample were also performed to rule out genomic DNA contamination. Following RT-PCR, 5 μl of each 25 μL reaction was run on a 3% agarose gel to assess the efficacy of the reverse transcription and amplification. The samples were then prepared for Pyrosequencing according to the standard recommendations for use with the PyroMark Q96 ID sequencer. The following primers were used for *Smcx* RT-PCR and Pyrosequencing: forward AGTCCAGGGCTGCTGCAGT; reverse, 5’-biotin GGAGCCAGCGTGCGTTTTA; sequencing, CAGGTGGAACAGGCG.

### Expression Plasmids and Lentiviral Transduction in ESCs

*mSmcx*, *hSMCX*, and *hSMCY* were cloned from cDNA libraries into the Gateway Entry vector (pENTR/D-TOPO, Invitrogen). The *SMCX* mutant H514A cDNA was generated by PCR-based mutagenesis. The sequence of all cDNAs was verified via Sanger sequencing. To generate the plasmids for stable overexpression in *X*^ΔTsix^*Y* male ESCs, the cDNAs were cloned into an in-house generated *PGK* promoter-driven Strep-tag II lentiviral vector by Gateway LR recombination. To generate doxycycline inducible constructs used in Fig. 3A-B, the cDNAs were cloned into a *PGK* promoter-driven HA-tag lentiviral vector pLIX-402 (Addgene plasmid # 41394) by Gateway LR recombination. For production of lentivirus, 50-70% confluent HEK293T cells in a 10-cm tissue culture plate were co-transfected with 3.33 μg lentiviral construct, 2.5 μg psPAX2 packaging plasmid and 1μg pMD2.G envelope plasmid using TransIT-293 (Mirus). Lentiviral particles were collected after 48 hrs of post transfection. The lentivral particles were then concentrated using LentiX (Clontech #631231). Concentrated lentivirus was transduced to ESCs with 10ug/ml Polybrene (Millipore, #TR-1003-G). After 72 hrs, the transduced cells were selected with 3ug/ml Puromycin (Sigma, #P8833-25MG) and subcloned to generate clonal cell lines. For inducible overexpression, cells were treated with 2ug/ml doxycycline (Sigma) for 3 days. Media were changed every 24 hours with fresh doxycycline.

### ChIP-Seq

ChIP was performed as previously described^55^, with a minor modification, where we sonicated the chromatin using truChIP sonicator (Covaris). Two independent *X*^ΔTsix^*Y*, *X*^ΔTsix;ΔSmcx^*Y*, or *Tg-mSmcx*;*X*^ΔTsix^*Y* ESC lines were differentiated into EpiLCs for all ChIP-Seq experiments with the exception that SMCX ChIP-Seq was performed on one *X*^ΔTsix;ΔSmcx^*Y* EpiLCs sample. To achieve optimal inter-sample normalization of ChIP efficiency, *Drosophila* chromatin, which contains fly-specific histone variant H2Av, was spiked in and subsequently immunoprecipitated with an anti-H2Av antibody (Active Motif) according to the manufacturers instruction. Two to five million EpiLCs were lysed and 3 μg of H3K4me2 antibody (Abcam #7766) or 10 μg of an in-house anti-SMCX antibody^55^ was used in individual ChIP reactions. ChIP-Seq libraries were sequenced on the Illumina HiSeq2500 platform to generate 50 bp single-end reads. Raw reads were demultiplexed and filtered according to the standard Illumina analysis pipeline. Reads from sequencing libraries were then mapped to the mouse (mm9) and fruit fly (dm6) genome assemblies using Bowtie^56^ allowing up to 2 mismatches. Only uniquely-mapped reads were used for analysis. Replicate BAM files were merged and reads were extended by 180 bp and converted to BED files. Basic sequencing statistics of all samples are presented in Methods Table 2.

For H3K4me2 analysis, *Drosophila*-normalized read coverage across the different genotypes was calculated over 1-kb bins across the mouse genome. Each 1-kb bin was normalized to the merged input samples. For SMCX, peaks were called using MACS2 software (version 2.1.0.20140616)^57^ using input bam files for normalization, with filters for a q-value < 0.1 and a fold enrichment greater than 1. This yielded 98 peaks on the X-chromosome for *X*^ΔTsix^*Y* samples and only 1 peak for the *X*^ΔTsix;ΔSmcx^*Y* sample. DNase I hypersensitive sites (DHS) were defined using MACS2 (q < 0.05) from the ENCODE mouse ESC data set^58^. Promoters were defined as ±1 kb from annotated transcription start sites (TSS) of the mm9 assembly. To find SMCX peaks, and subsequently H3K4me2 density, at DHS or promoters, we used the Bedtools intersect command to select SMCX peaks in which 10% of the SMCX peak length overlapped the promoters or DHS. For visualization in the Integrated Genome Viewer, bigwig files were generated with coverage normalized using the number of mapped reads to the *Drosophila* genome (reads mapped per reference genome per million reads)^59,60^.

### ChIP-qPCR

ChIP was performed as above. Two to five million cells were lysed and 3 μg of H3K4me1 antibody (Abcam #8895) or 3 μg of H3K4me2 antibody (Abcam #7766) or 3 μg of H3K4me3 antibody (Abcam #8580) or 3 μg of H3K27ac antibody (Active motif #39135) or 3 μg of rabbit IgG antibody (Jackson Immuno Research #011-000-003) was used in individual ChIP reactions. DNA was purified by phenol/chloroform extraction and eluted using QIAquick purification kit (QIAGEN#28104). All qPCR reactions were performed using SYBR Green-based relative quantification method on an Eppendorf Realplex Mastercycler.

### Fluorescent Western blotting

The Odyssey CLx system (LI-COR Biosciences, Lincoln, NE, USA) was used for fluorescent Western blotting. Protein lysates were made in RIPA buffer (50mM Tris-HCl, 1% NP40, 0.25% Na-deoxycholate, 150mM NaCl) with PMSF (Sigma, #P7626) and protease inhibitor cocktail (Roche, #11873580001). Lysates were mixed with Laemmli sample buffer and boiled for 10 min prior to loading on Polyacrylamide gels. Resolved proteins were transferred onto PVDF membranes (Immobilon-P, Millipore) overnight. Membranes were then blocked in 5% BSA (Sigma, #A7906) for 1 hr at room temperature, and incubated in the anti-Strep-Tag II (GenScript, #A01732-100) and anti-Actin (Sigma, #A5060) primary antibodies together overnight at 4°C. Appropriate Li-COR IRDye series (680LT/800LT) secondary antibodies were used. Blots were scanned using the Odyssey CLx Imager (Li-COR Biosciences) following the manufacturer’s instruction. Results were analyzed using Odyssey Li-COR software. Transgenic SMCX levels were normalized to ACTIN in each blot.

### Microscopy

Images of all stained samples were captured using a Nikon Eclipse TiE inverted microscope with a Photometrics CCD camera. The images were analyzed after deconvolution using NIS-Elements software. All images were processed uniformly.

### SMCX/SMCY Sequence Analysis

Amino acid sequences were obtained from the Ensembl database and from Cortez *et al*., 2014^61^. Multiple sequence alignment was performed using MSAViewer^62^. MSAViewer tree tool was used to create a rooted phylogenetic tree with branch lengths according to the Neighbor Joining algorithm. Finally, the alignment was analyzed to find amino acids conserved across mammalian (human, mouse, and elephant) SMCX, but not in mammalian SMCY sequences nor in the orthologous chicken KDM5A sequence. These sites were visualized within the domain structure of human SMCX using PROSITE (http://prosite.expasy.org/), an available ExPASy bioinformatic resource tool.

## Acknowledgments

We thank members of the Kalantry and Iwase laboratories for discussions and critical review of the manuscript; Jacob Mueller for critically evaluating the manuscript; Angela Andersen of Pickersgill and Andersen, Life Science Editors (lifescienceeditors.com/), for editing services. We thank Michael Hinten for designing the *Smcx* Pyrosequencing assay; and, Paul Ginart and Arjun Raj for help with designing of the allele-specific Xist RNA FISH assay; and Saurabh Agarwal for technical assistance for genomic analyses. We acknowledge the services of the University of Michigan Sequencing Core Facility, supported in part by the University of Michigan Comprehensive Cancer Center. This work was funded by NIH National Research Service Awards 5-T32-GM07544 (University of Michigan Predoctoral Genetics Training Program; to E.M. and C.N.V.), T32-HD079342 (University of Michigan Predoctoral Career Training in the Reproductive Sciences Program; to E.M., C.N.V., and R.S.P), 1F31HD080280-01 (to E.M.), and a National Science Foundation Graduate Research Fellowship DGE-1256360 (to P.M.G.), a Rackham Predoctoral Fellowship from the University of Michigan (to E.M.), an NIH Director’s New Innovator Award (DP2-OD-008646) (to S.K.), a March of Dimes Basil O’Connor Starter Scholar Research Award (5-FY12-119) (to S.K.), NIH NINDS Award (R01NS089896) (to S.I.), Farrehi Family Foundation Grant (to S.I.), a Reproductive Science Program Pilot Grant (to S.K. and S.I.), and the University of Michigan Endowment for Basic Sciences (S.K. and S.I.). Sequencing data generated for this study have been submitted to the NCBI Gene Expression Omnibus (GEO; <u>http://www.ncbi.nlm.nih.gov/geo/</u>) under accession number GSE96740.

## Author Contributions

S.G., S.I., and S.K. conceived the study and designed the experiments. S.I. generated *Smcx*^fl^ mouse strain. S.G. bred *Smcx* mice, derived, characterized, transduced, and analyzed ESC, EpiLC, and EpiSC lines. E.M. performed dissections and imaging of post-implantation embryos. C.N.V. prepared RNA-seq libraries and performed RNA-seq analyses, and assisted with the preparation of SMCX/SMCY constructs. Y.M.N prepared ChIP-Seq libraries. R.S.P. and P.M.G. performed ChIP-Seq analyses. All authors contributed to the writing and editing of the manuscript.

## Availability of Data and Material

Sequencing data generated for this study have been submitted to the NCBI Gene Expression Omnibus (GEO; <u>http://www.ncbi.nlm.nih.gov/geo/</u>) under accession number GSE96740. All cell lines generated in this study are available upon request.

